# A workflow using diaPASEF global quantitative proteomic analysis reveals extracellular vesicle biomarker candidates for non-invasive diagnostics in non-small cell lung cancer

**DOI:** 10.1101/2025.10.26.684684

**Authors:** Manan Vij, Patrick Kurnia, Lyssa Dimapanat, Rajesh K. Soni, Alex J. Rai

## Abstract

Lung cancer is the second most diagnosed cancer in the world. Non-small cell lung cancer is the most common type of lung cancer in the United States. Tissue biopsy is the gold standard for detecting lung cancer but is highly invasive as it necessitates the extraction of a sample of tissue for histologic analysis. It also carries risks of bleeding and/or infection and is inconvenient from a patient perspective. The development of a minimally invasive test, such as one utilizing a blood or urine sample, and capable of providing accurate results for lung cancer detection and/or subtyping, would significantly enhance the clinical landscape and streamline patient care. In this study we utilize A549 and H1299 human lung cancer cell lines, differing in cell type, location within the lung, and genetic composition (Kras & p53 status), and employ diaPASEF for global quantitative proteomic analysis. We demonstrate that extracellular vesicle protein content provides enhanced resolution to differentiate between these two cell lines relative to protein lysate content and reveals alterations in signaling. Protein clusters are identified showing enrichment for distinct biological processes representing multiple gene ontology categories unique to each lung cancer subtype-oxidative phosphorylation, apical junction, and epithelial-mesenchymal transition. We subsequently delineate a short list of urine-detectable protein candidates that is prognostic in a second cohort of lung cancer patients. This list of protein candidates may be useful for the development of a non-invasive test to distinguish between these two subtypes of human lung cancer.

## Introduction

Lung cancer is the second most diagnosed lung cancer in the world and remains the leading cause of cancer deaths across the world [1]. Non-small cell lung cancer (NSCLC) is the most common type of lung cancer within the US, causing more deaths than colorectal, breast and prostate cancers combined, and includes a variety of different lung cancers, including adenocarcinoma, squamous cell carcinoma, and large cell carcinoma [2, 3]. Despite a variety of imaging technologies being used to identify lesions suspected as cancerous, the ability of current imaging techniques to effectively diagnose lung cancer is poor. As a result, tissue biopsy procedures play a critical role in the clinical pathway for lung cancer diagnosis [3]. While tissue biopsy is the gold standard for detecting lung cancer, it is highly invasive as it necessitates the extraction of a sample of tissue for histological analysis. It also carries risks of bleeding and/or infection and is inconvenient from a patient perspective. Furthermore, tissue biopsy is a feasible clinical option only when a suspected cancerous mass becomes detectable [4].

In contrast, liquid biopsy offers a promising paradigm shift in cancer diagnostics by leveraging the analysis of circulating tumor components in bodily fluids [5]. Liquid biopsy holds the potential for early cancer detection, as it is not hindered by the constraints that govern tissue biopsy procedures. Utilizing minimally invasive tests such as blood or urine tests present an opportunity to revolutionize the landscape of lung cancer diagnosis [5]. Such a test has the potential to streamline diagnostic processes, facilitate early intervention, and improve patient outcomes. Thus, the development of a minimally invasive test, such as one utilizing a blood or urine sample, capable of providing accurate results for lung cancer detection and/or subtyping, would significantly enhance the clinical landscape and streamline patient care.

In this study we utilize A549 and H1299 human lung cancer cell lines. The A549 tumor-cell line, initiated from a human alveolar cell carcinoma, expresses characteristic features of alveolar type II cells, including synthesis of phospholipids, cytoplasmic lamellar bodies, and apical microvilli [6, 7]. It also permits in vitro analysis of human surfactant synthesis and secretion [7]. H1299 cell line is established from a lymph node metastasis of the lung from a 43-year-old white male patient with carcinoma. While both these cell lines are NSCLC-derived, they differ in genetic composition of p53 and KRAS expression, two proteins whose downstream signaling pathways are crucial for developing effective cancer therapies [8]. We demonstrate herein that the extracellular vesicle protein content derived from these two cell lines provides sufficient predictive power to differentiate between these two cell lines and identify differentially enriched biological processes and associated proteins for each lung cancer subtype. Specifically, H1299 cell lines show enrichment in apical junction and epithelial-mesenchymal transition ontologies, while A549 cell lines show enrichment in proteins related to oxidative phosphorylation. In addition, we identify proteins that are detectable in urine and query a second cohort of lung cancer patients to identify those proteins that have prognostic value. This short list of suitable biomarker candidates may be useful for the development of a non-invasive urine-based laboratory test to detect or distinguish the two lung cancer subtypes using a combination of clinically relevant biomarkers.

## Materials & Methods

### Cell lines used-

Lung cancer cell lines, A549 and H1299, in addition to two control cell lines (SCBO-23, bladder cancer, and HeLa, cervical cancer) were used for comparative analysis. All cell lines were purchased from ATCC (Maryland/Virgina, USA), except for SCBO-23 cell line which was a gift from laboratory of Michael Shen, PhD, at Columbia University Irving Medical Center.

### Isolation of EVs and comprehensive proteomics analysis-

Extracellular vesicles were isolated from cell lines kept in serum-free media for 24 hours, as previously described [9–10]. Briefly, media was collected and subjected to serial centrifugation steps at 3K x g, 20K x g, and 100K x g, to remove dead cells and cell debris, large microvesicles and protein aggregates, and exosomes, respectively [9]. The 100K pellet was further analyzed using global quantitative proteomic analysis, as described below.

Global quantitative proteomic analysis-For the global quantitative proteomic analysis of extracellular vesicles, diaPASEF [11] based proteomics was employed. In brief, cells were lysed in SDC buffer [11] (1% SDC, and 100 mM Tris-HCl, pH 8.5) and boiled for 15 min at 60°C, 1000 rpm. Protein reduction and alkylation of cysteines was performed with 10 mM TCEP and 40 mM CAA at 45°C for 10 min followed by sonication in a water bath, cooled down to room temperature. Protein digestion was processed for overnight by adding LysC / trypsin mix in a 1:50 ratio (µg of enzyme to µg of protein) at 37° C and 1400 rpm. Peptides were acidified by adding 1% TFA, vortexed, and subjected to StageTip clean-up via SDB-RPS [12] and dried in a speed-vac. Peptides were resuspended in 10 μl of LC buffer (3% ACN/0.1% FA). Peptide concentrations were determined using NanoDrop, and 200 ng of each sample was used for diaPASEF analysis on timsTOFPro.

### Liquid chromatography with tandem mass spectrometry (LC-MS/MS)-

Peptides were separated within 87 min at a flow rate of 300 nl/min on a reversed-phase C18 column with an integrated CaptiveSpray Emitter (25 cm x 75µm, 1.6 µm, IonOpticks). Mobile phases A and B were with 0.1% formic acid in water and 0.1% formic acid in ACN. The fraction of B was linearly increased from 2 to 23% within 60 min, followed by an increase to 35% within 7 min and a further increase to 90% before re-equilibration. The timsTOF Pro was operated in in diaPASEF mode [11] and data was acquired at defined 32 × 26 Th isolation windows from m/z 400 to 1,200. The collision energy was ramped linearly as a function of the mobility from 59 eV at 1/K0=1.6 Vs cm^-2^ to 20 eV at 1/K0=0.6 Vs cm^-2^.

The acquired diaPASEF raw files were analyzed using the Spectronaut software platform (Biognosys) in library-free mode. Default parameters were used except that decoy generation was set to “mutated.” Data were searched against the human reference proteome downloaded from UniProt. The false discovery rate (FDR) was controlled at 1% at both the peptide precursor and protein levels using the mProphet scoring algorithm [13].

### Bioinformatics analysis-

Cell lysate and extracellular vesicle protein-expression data were analyzed using ProteoDisco 2.0, a custom-designed app for analysis of proteomics data in R. A manuscript describing all functionality for Proteodisco 2.0 is under preparation (A.J. Rai). Proteomics data was scaled using quantile normalization to mitigate external, non-biological differences between the samples. Differential protein expression testing was performed using Student’s t-test to determine significant differentially expressed proteins among the samples. Principal Component Analysis (PCA) of extracellular vesicle and lysate protein samples was used to visualize the expression matrix in two and three dimensions. The first and second principal components were selected to generate the PCA plots.

Survival analysis was conducted employing the mRNA lung cancer specific module in Kaplan-Meier Plotter [14]. Queries were run using default parameters, to associate survival outcomes with expression level of candidate biomarkers to generate survival curves. The demographic data reported (schematic in Figure 5) was obtained from Gyorffy B. et al [14].

## Results

### Proteomic Characterization of A549 and H1299 Cell Lines

#### Extracellular vesicle protein content provides better separation to distinguish between A549 and H1299 NSCLC relative to cell lysate protein content-

Figure 1 depicts Venn diagrams corresponding to the number of proteins detected in cell culture lysates and EVs isolated from the culture media of the corresponding cell lines. Spearman correlation coefficient matrix across all samples is also depicted.

**Figure 1.**
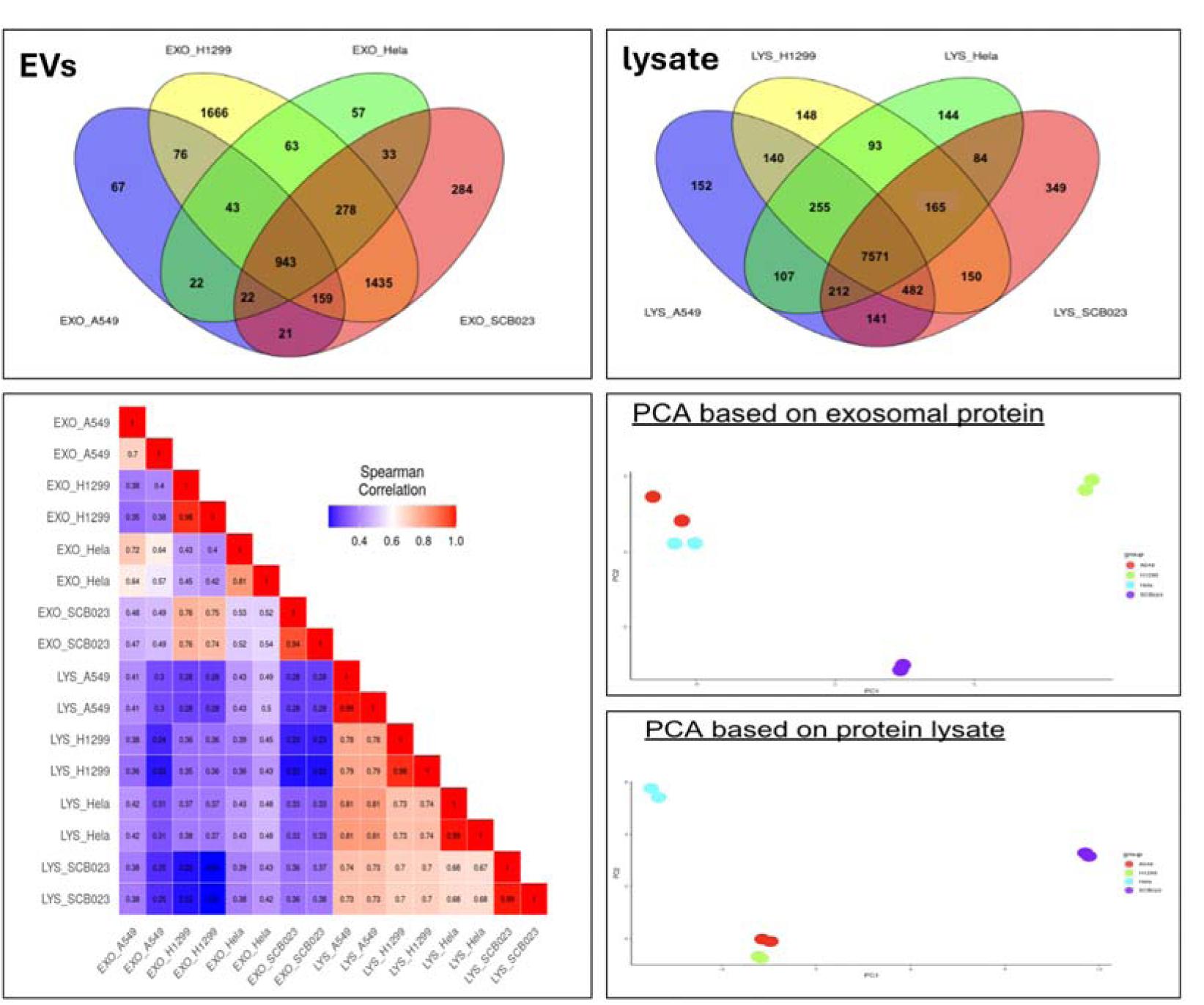
Venn diagram indicating total number of proteins detected in all samples, extracellular vesicles (left) and lysates (right). Spearman correlation matrix for all samples. 2D PCA plot for EV (top) and lysate (bottom) samples; each dot is a replicate.

Two-dimensional Principal Component Analysis (PCA) plots are also shown. Figure 2 depicts all Volcano plots showing binary comparisons of the various cell lines.

**Figure 2:**
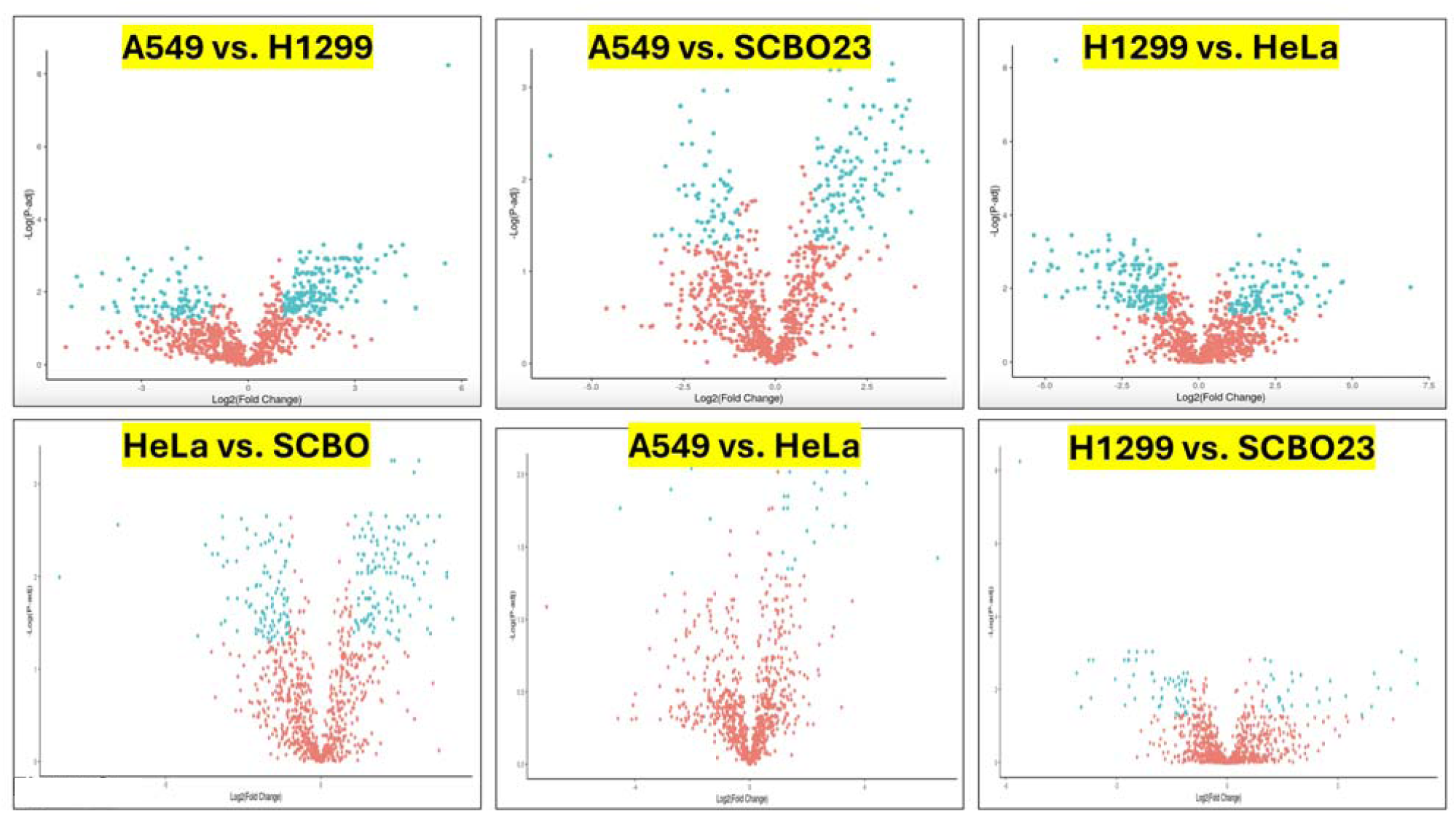
Volcano plots for all binary comparisons. Proteins exhibiting statistically significant differences and >2x fold change are indicated in blue color.

#### Functional enrichment and affected signaling pathways-

Results from GSEA and gene ontology analysis are shown in Figure 3. Heatmaps depicting the relative levels of specific proteins for each ontology are also shown. A549 cell lines showed enrichment in proteins involved in Oxidative Phosphorylation, while H1299 cell lines were enriched for Epithelial-Mesenchymal Transition and Apical Junction related pathways. Figure 3 also shows the GSEA enrichment score plot for each of the statistically significant pathways identified by gene ontology analysis.

**Figure 3:**
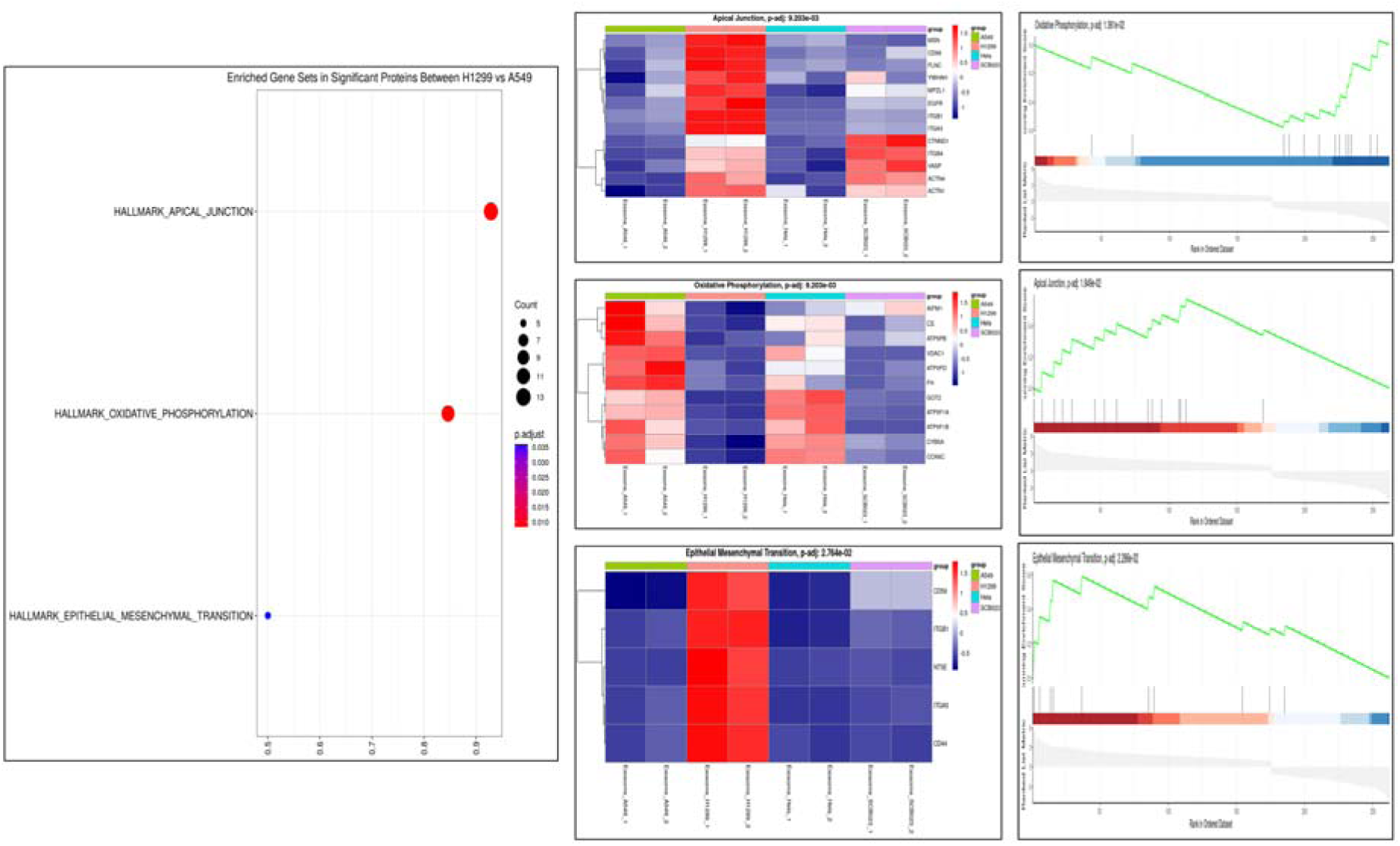
Gene ontology and heatmap including specific proteins for each of three significant cancer ontologies are depicted. Individual proteins involved in apical junction pathways, oxidative phosphorylation, and epithelial mesenchymal transition pathways are depicted. Gene set enrichment analysis results for the three pathways are shown.

Figure 4 depicts a chord diagram with the top few proteins of each of the three significant pathways. Two proteins, EGFR and ITGA5, map to more than one category, and a total of 14 proteins depicted in the chord diagram constitute a biomarker signature that maps to the three affected pathways distinguishing the A549 and H1299 lung cancer cell lines.

**Figure 4.**
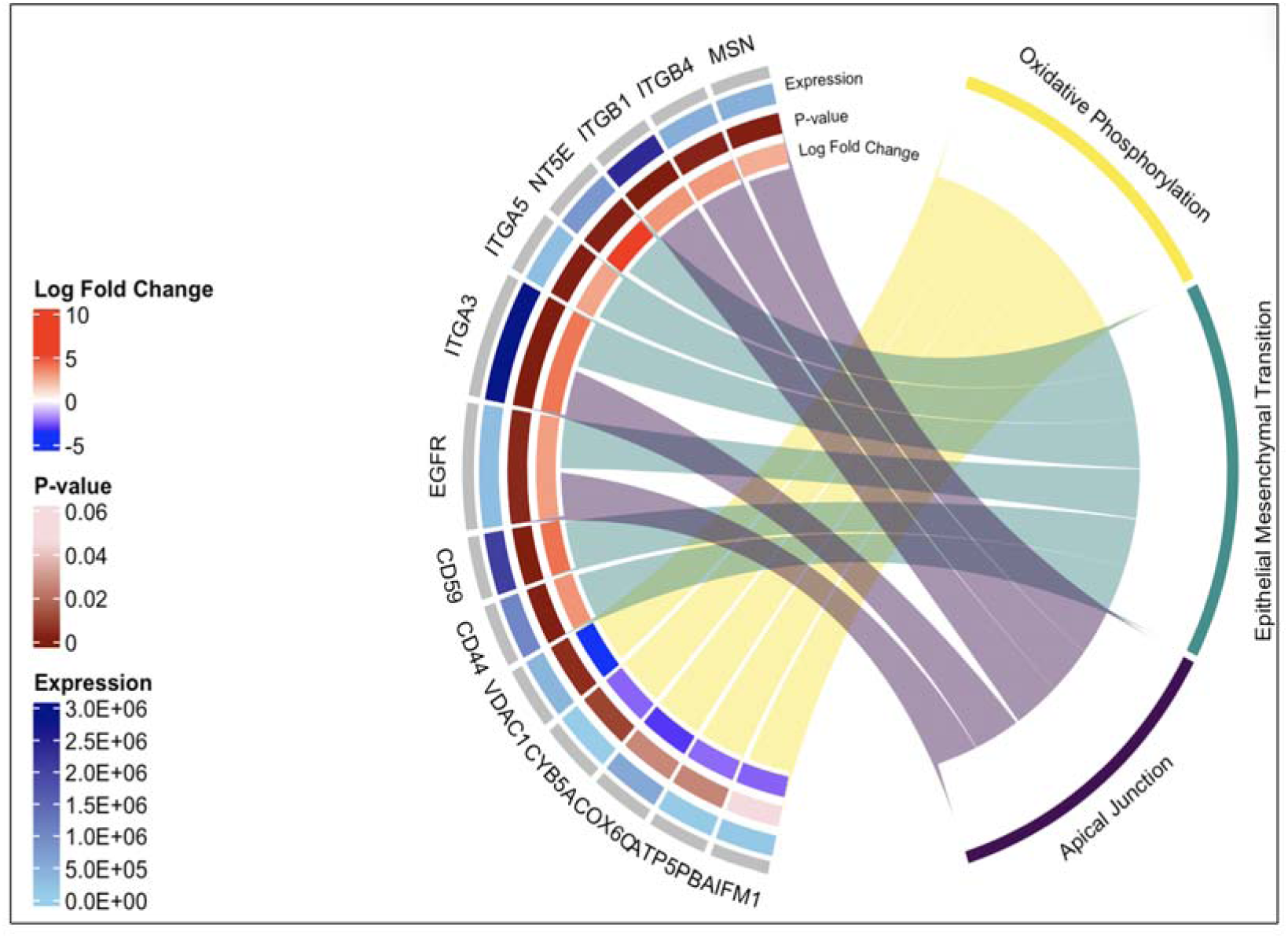
Chord Diagram showing top 14 proteins associated with each of the identified cancer pathway from GSEA.

**Figure 5:**
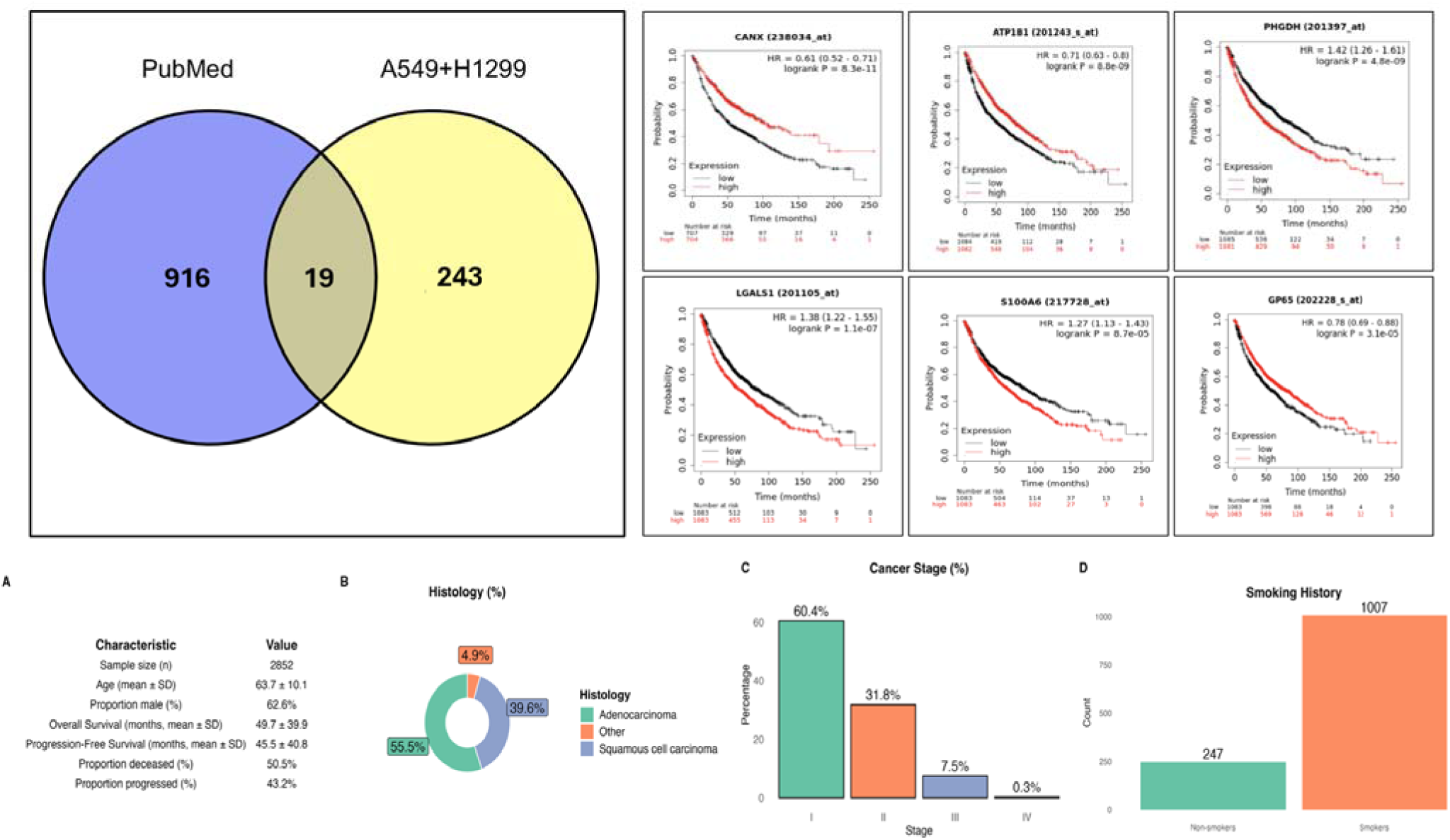
Venn diagram depicting overlap of urine-detectable protein biomarkers of lung cancer (PubMed query) and significant proteins from current analysis of A549 and H1299 cell lines. Of 19 significant candidates, top 6 most statistically significant candidates are depicted (right), showing the Kaplan-Meier (K-M) survival analysis based on RNA expression levels of candidates in all lung cancer cases using default parameters. Bottom, demographic data for cases included in K-M plots.

### Urine-detectable proteins, assessment in second cohort, and pruning of Candidate biomarker list

#### Literature Review-

In an effort to prune our list of 243 candidate biomarkers and prioritize them for further validation, we performed a PubMed query using the following search terms: “lung cancer”, “urine”, “protein biomarker.” The search was restricted to the last 10 years and resulted in a total of 110 articles (Table 1). Manual review of the papers identified 916 protein candidates (proteins having aliases were included as separate entries). Figure 5 shows a Venn diagram depicting the overlap between protein biomarkers identified from EV proteomics analysis comparison of H1299 and A549 cell lines and the literature search; a total of 19 proteins were common to both.

**Table 1:**
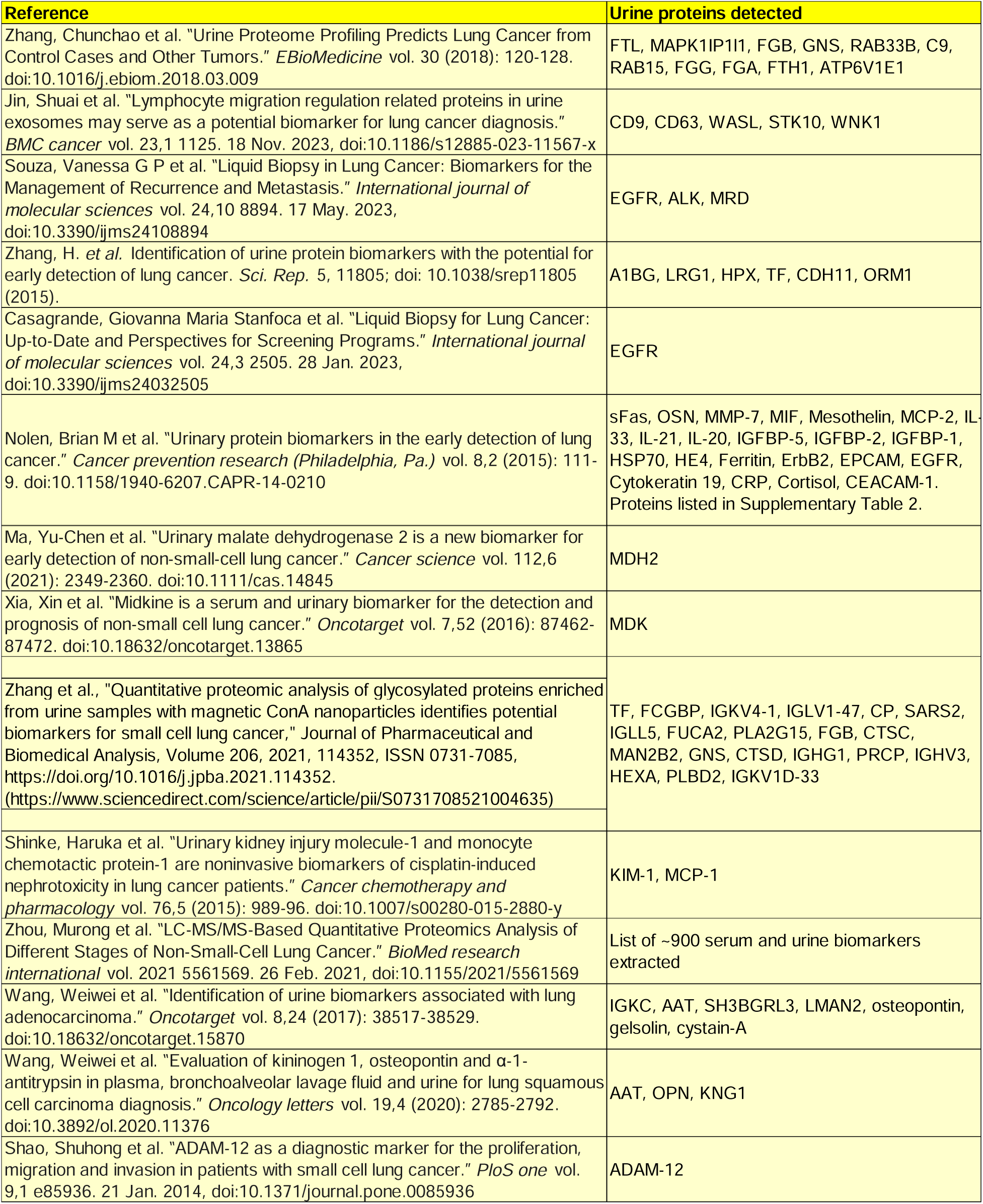
PubMed search results tabulation.

#### Validation in separate cohort-

These nineteen proteins were queried in a separate cohort of lung cancer patients to test their association with survival outcome. This analysis was conducted using K-M Plotter [13]. Sixteen proteins retain prognostic significance. Table 2 shows the sixteen candidate biomarkers and the P-value and hazard ratios corresponding to these proteins in the cohort (details on cohort demographics are also depicted in Figure 5).

**Table 2:** Nineteen candidate proteins overlapping between comparison of A549 & H1299 cell lines and PubMed protein search results.

| A549 |  |  | H1299 |  |  |
| --- | --- | --- | --- | --- | --- |
| Protein | Logrank p-value | Hazard Ratio | Protein | Logrank p-value | Hazard Ratio |
| S100A6 | 8.70E-05 | 1.27 | PHGDH | 4.80E-09 | 1.42 |
| CANX | 8.30E-11 | 0.61 | EGFR | 0.0027 | 0.8 |
|  |  |  | ATP1B1 | 8.80E-09 | 0.71 |
|  |  |  | LGALS1 | 1.10E-07 | 1.38 |
|  |  |  | RALB | 0.01 | 0.86 |
|  |  |  | EPHA2 | 0.017 | 0.86 |
|  |  |  | RRAS2 | 0.00022 | 0.8 |
|  |  |  | RAC1 | 0.0185 | 0.87 |
|  |  |  | TPM4 | 0.00256 | 0.79 |
|  |  |  | CXADR | 0.0014 | 0.79 |
|  |  |  | SH3BGRL3 | 0.00478 | 1.19 |
|  |  |  | TLN1 | 0.0188 | 1.15 |
|  |  |  | NPTN (GP65) | 0.00003 | 0.78 |
|  |  |  | CD9 | 0.01505 | 1.09 |

## Discussion

The present investigation addresses the urgent need for a less invasive diagnostic test for non-small cell lung cancer (NSCLC), the predominant lung cancer subtype within the United States and the second most common globally. The conventional gold standard, tissue biopsy, is associated with considerable invasiveness, concomitant risks of bleeding and infection, and patient inconvenience. Consequently, the identification of a minimally invasive alternative, specifically one utilizing blood or urine, would refine the clinical paradigm and optimize patient care. In this study, A549 and H1299 human lung cancer cell lines, distinguished by cell type, lung location, and genetic composition (Kras and p53 status), were used. A549 cells, originating from type II alveolar epithelium, exhibit a KRAS mutation, while H1299 cells, derived from bronchiolar epithelial cells, manifest a p53 deletion mutation. Our analysis highlights the potential of extracellular vesicle protein cargo from these cell lines to predict and differentiate the two subtypes of lung cancer and provides a short list of biomarker candidates to potentially carry forward for the development of a non-invasive laboratory test utilizing urine samples.

For the lysates, over 7500 proteins are common to all samples with <2% being unique to any of the cell lines. In the EV preparations, 943 proteins were common to all samples, with ∼1500-4000 proteins detected in each sample.

The largest differences in protein content were seen between the lysate and EV preparations, while the lysates of the various cell lines were more similar to each other. The EV samples showed an intermediate correlation-some were more closely correlated than others. The similarity in protein composition of the lysates likely reflects the large number of housekeeping proteins common to all cells. This was apparent from the correlation coefficient matrix.

PCA plots of EVs demonstrated a high degree of discrimination between A549 and H1299 lung cancer cell lines. Interestingly, results revealed similar EV profiles between A549 and Hela cell lines, in agreement with the correlation analysis. Lysate PCA graph revealed that the two lung cancer cell lines had similar protein profiles based on cell lysate proteins. Replicates showed close clustering indicating good reproducibility among the replicates. Thus, our analysis demonstrated that EV protein content was better able to distinguish A549 and H1299 lung cancer subtypes relative to the protein lysate from the cell lines.

Our analysis sheds light on the potential utility of EV protein content to differentiate between two lung cancer cell types. The observed disparities in protein overlap and the low correlation in protein expression levels between the two lung cancer cell lines signify that EV proteins effectively capture divergent biological characteristics. This interpretation is reinforced by the identification of distinct biological pathways enriched in the A549 and H1299 cell lines, underscoring the nuanced nature of their respective protein profiles. The manifestation of these disparate pathways, attributed to the differential expression of proteins between the two cell lines, reveals unique signaling in each lung cancer subtype.

Comparison of the A549 and H1299 cell lines (Figure 2), revealed MT2A, NT5E, B2M and CD59 as among the top 25 proteins characteristic of A549 that are also detectable in urine. Similarly, S100A6, FABP4, KRT7, ALDH1A1, and KRT13 were identified as urine detectable biomarkers characteristic of H1299. Comparisons of the two lung cancer cell lines with SCB023 and HeLa cell lines were performed as controls for the current analysis and will not be discussed herein.

This study provides proteomic characterization of these differences and uncovers potential biomarkers that can be used to distinguish the cell lines and to uncover biological pathways activated within each subtype. Other recent studies have also identified potential biomarkers to differentiate lung cancers from other cancer types [15–17]. However, lung cancers themselves are a heterogenous group of cancers. Our analysis extends the existing studies by providing biomarkers to distinguish between two common subtypes of lung cancer that could potentially serve as targets in non-invasive clinical lab testing for lung cancer subtyping. These urine-detectable proteins include MT2A, NT5E, B2M and CD59 for the A549 cell line, and S100A6, FABP4, KRT7, ALDH1A1, and KRT13 for the H1299 cell line. Our candidate proteins contribute novel biomarker candidates to improve diagnostic strategies for NSCLC.

Our analysis identified 19 protein candidates that were previously detected in urine samples from human lung cancer patients and subsequent validation in a second cohort of lung cancer proteins demonstrated prognostic significance for 16 of these proteins-two in A549 lung cancer cells (S100A6, and CANX) and fourteen from H1299 lung cancer cells (PHGDH, EGFR, ATP1B1, LGALS1, RALB, CD9, EPHA2, RRAS2, RAC1, TPM4, CXADR, SH3BGRL3, TLN1, and NPTN). Considering these previous results, we suggest that, after appropriate validation, these protein biomarkers may be useful for non-invasively subtyping human lung cancer, and after further validation, could obviate the need for tissue biopsy.

Our study also assessed these biomarkers for their application as prognostic biomarkers for non-small cell lung cancer (NSCLC). S100A6 and CANX protein urine biomarkers from A549 cells were found to be strongly correlated with NSCLC patient survival outcomes, with high S100A6 expression correlated with poor prognosis and high CANX correlated with improved prognosis. Similarly, among H1299 cells, high expression of PHGDH, LGALS1, SH3BGRL3, and TLN1 were associated with worse outcomes, while high expression of EGFR, ATP1B1, RALB, EPHA2, RRAS2, TPM4, CXADR, GP65, and RAC1 were associated with improved survival. This group of proteins underscores the depth and breadth of our study, showcasing numerous potential targets useful for refining prognostication and guiding therapeutic interventions in non-small cell lung cancer patients. These proteins not only hold significant diagnostic and prognostic value for both lung cancer subtypes, but also serve as detectable markers, useful for guiding clinical decision-making in lung cancer management.

There are caveats to our current findings and conclusions. Firstly, the clinical utility of these biomarkers needs to be further validated in larger patient cohorts from different sites. Secondly, it should be noted that the analysis using the validation cohort herein is based on RNA levels from lung cancer tissue samples. Changes in RNA levels do not necessarily reflect altered biomarker protein levels that are detectable in urine. Finally, while our study identifies multiple versatile biomarkers for lung cancer subtypes exemplified by the two lung cancer cell lines, the study is limited by experimental validation of these protein biomarkers in vivo using other methods. Nevertheless, this current analysis will inform future experiments that focus on comprehensive experimental validation to confirm the detection/quantitation of the identified protein biomarkers in both lung cancer cell lines and in patient-derived urine samples. Through such analyses, we aim to demonstrate the utility and reliability of these biomarkers for clinical applications in lung cancer detection, subtyping, prognosis, and/or treatment stratification.

